# Ebola outbreak brings to light an unforeseen impact of tsetse control on sleeping sickness transmission in Guinea

**DOI:** 10.1101/202762

**Authors:** Moïse Kagabadouno, Oumou Camara, Mamadou Camara, Hamidou Ilboudo, Mariame Camara, Jean-Baptiste Rayaisse, Abdoulaye Diaby, Balla Traoré, Mamadou Leno, Fabrice Courtin, Vincent Jamonneau, Philippe Solano, Bruno Bucheton

**Affiliations:** Programme National de Lutte contre la Trypanosomiase Humaine Africaine (PNLTHA), Ministère de la Santé, République de Guinée; Centre International de Recherche Développement sur l’Elevage en zone Subhumide (CIRDES), Burkina Faso; Direction Préfectorale de la Santé de Boffa, Ministère de la Santé, République de Guinée; INTERTRYP, IRD, CIRAD, Université Montpellier, Institut de Recherche pour le Développement, 34398 Montpellier, France

## Abstract

In addition to the thousands of deaths due the unprecedented ebola outbreak that stroke West Africa (2014-2016), national health systems in affected countries were deeply challenged impacting a number of diseases control programs. Here we describe the case of Human African Trypanosomiasis (HAT), a deadly neglected tropical disease due to a trypanosome transmitted by tsetse flies for which no vaccine nor chemoprophylaxis exists. Data are presented for the disease focus of Boffa in Guinea where a pilot elimination project combining medical screening and vector control was launched in 2012. During ebola, HAT active screening activities were postponed and passive surveillance also was progressively impaired. However, tsetse control using small insecticide impregnated targets could be maintained. The over two years disruption of screening activities led to a dramatic increase of HAT prevalence, from 0.7% in 2013 (21/2885) to 2% (69/3448) in 2016, reaching epidemic levels (>5%) in some villages. In deep contrast, control levels reached in 2013 (0.1%; 7/6564) were maintained in areas covered with impregnated targets as no cases were found in 2016 (0/799). In Boffa, ebola has thus incidentally provided a unique framework to assess the impact of current HAT control strategies. A first lesson is that the “screen and treat” strategy is fragile as rapid bursts of the disease may occur in case of disruption. A second lesson is that vector control reducing human-tsetse contacts, even implemented alone, is effective in providing a good level of protection against infection. This advocates for a greater attention being paid to the combination of tsetse control together with medical activities in aiming to reach the HAT elimination objective in Africa.

## Vector control and HAT elimination: an ongoing debate

Human African Trypanosomiasis (HAT) or sleeping sickness, is a deadly neglected tropical disease transmitted to humans by tsetse flies. Elimination as a public health problem was targeted for 2020^1^. Notwithstanding, major challenges are still to be faced regarding sustainability and transmission interruption^2^. The dominating screen and treat strategy, used to clear the human reservoir, is progressively switching to more cost-effective passive surveillance systems based on the recent availability of rapid diagnostic tests^3^. Long overlooked, tsetse control has only been re-introduced recently in a few HAT foci and debates are ongoing concerning its efficacy and cost effectiveness^4^. A pilot study, launched in the Boffa mangrove focus (Guinea) in 2012, has shown that reducing human exposure with tsetse attractive insecticide impregnated targets (tiny targets), significantly enhanced the impact of screening campaigns on transmission^5^. Indeed, in the Eastern part of the focus where tiny targets were deployed, a reduction by three of the HAT prevalence was observed in 2013 only one year after initial deployment with a very low incidence of new infections below the level of 0.1 %. The same approach has been also used with success in Chad^6^.

## A unique opportunity to assess the impact of tsetse control provided by the ebola outbreak

The unprecedented Ebola outbreak that stroke Guinea (2014-2015) had dramatic impacts affecting most health programs^7^. HAT control was deeply hampered as no active screenings could be led during this period. Passive detection with rapid tests in health centers was organized early in 2014, but was efficient only for several months^8^ and Dubreka, located a 100 km from Boffa, became rapidly the only functional site for diagnosis and treatment of HAT in Guinea. Remarkably, vector control initiated on the Eastern bank of the Rio Pongo River in the Boffa focus, could be maintained throughout the ebola period. This was thanks to a strong adhesion of the population who participated actively in the yearly renewal of tiny targets around their villages in 2014 and 2015 with almost half of the targets being replaced by the villagers themselves (Figure 1a,1b). Tsetse flies densities could thus be kept under control in the Eastern part of the Boffa focus with a greater than 80 % reduction level as compared to the pre-intervention period (Figure 1c). In the Boffa focus, ebola has thus unexpectedly provided a unique opportunity to assess the impact of vector control on transmission in the absence of medical screening.

**Figure 1:**
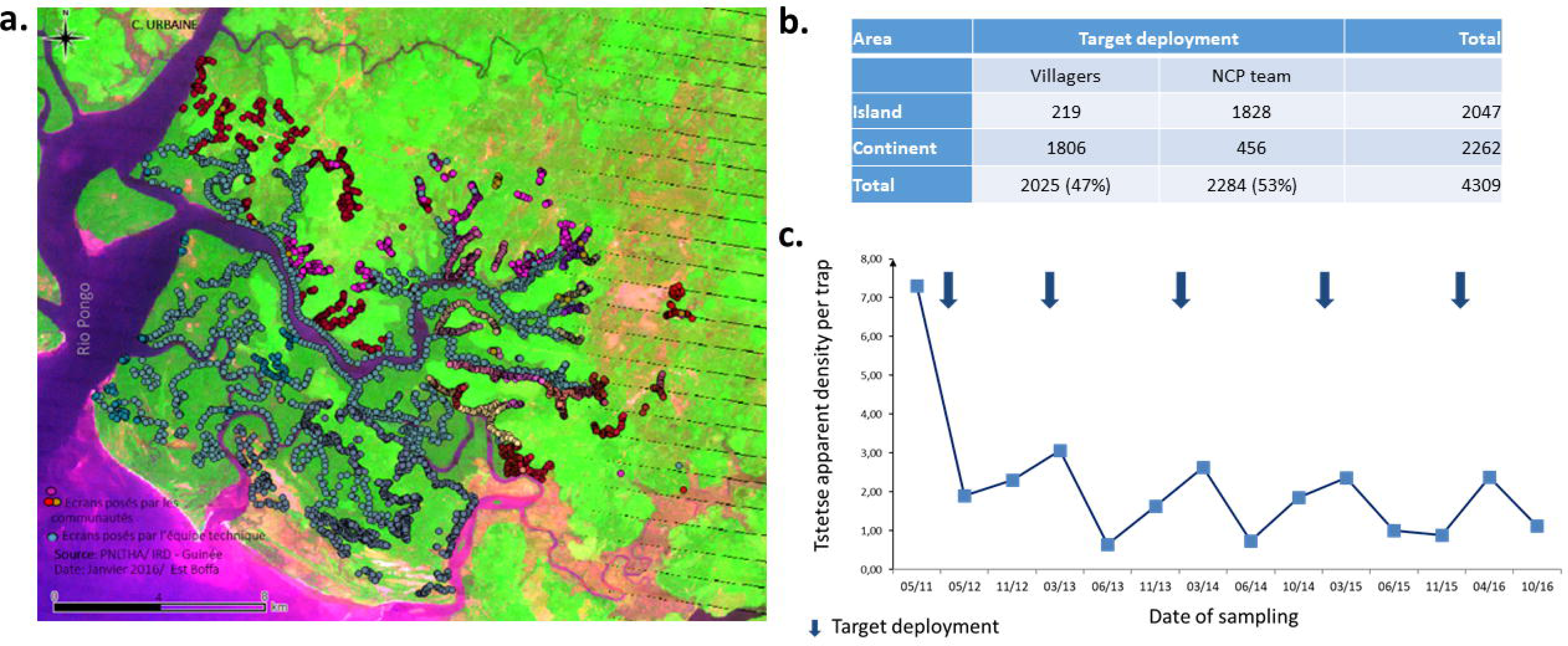
**a.** Geographic distribution of insecticide impregnated targets on the Eastern bank of the Rio Pongo River in the Boffa HAT focus after the deployment in January 2016; b. Number of tiny targets deployed in 2016 by the NCP team and by the villagers themselves; c. Tsetse densities were assessed by setting biconical traps for 48 hours at 47 geo-referenced sentinel sites; **d.** Pictures of target deployment targeted to reduc human-tsetse contacts, from right to left: assembly of tiny targets by the villagers on bamboo sticks; deployment of targets in mangrove channels by the NCP team; tiny targets at a pirogue jetty site, tiny targets deployed around a rice scheme, tiny target at a watering point, tiny targets at a river site used for washing activities by the villagers.

After Guinea was declared free of Ebola in 2016, two medical surveys could be implemented in the focus in May and October. Altogether, 79 HAT cases were diagnosed out of 4,247 persons screened. This represents a sharp increase of disease prevalence as compared to the pre-ebola period (from 0.7 to 2%), with villages reaching epidemic levels with up to 5% of prevalence. Strikingly all cases were detected in villages where no vector control activities had occurred (Figure 2a), on the Western part of the focus. In the eastern part where vector control had been implemented, no case could be detected. The relatively low number of individuals screened (799) in the vector control intervention area in the Eastern part of the focus, was due to the maximization of screening efforts toward most affected populations and a low participation of the population in this area where HAT does no longer appear as a major health problem. This low number of people screened precludes drawing firm conclusion on transmission interruption in this area, nevertheless the absence of HAT cases observed during this survey indicates that the pre-ebola control level in Boffa East was at least maintained (Figure 2b) since the last survey held in 2013 (0.1 % prevalence). In line with this statement is the fact that among the 22 HAT cases passively diagnosed from the Boffa focus during the ebola outbreak (2014-2015), only two came from villages where tiny targets were deployed and replaced each year since 2012.

**Figure 2:**
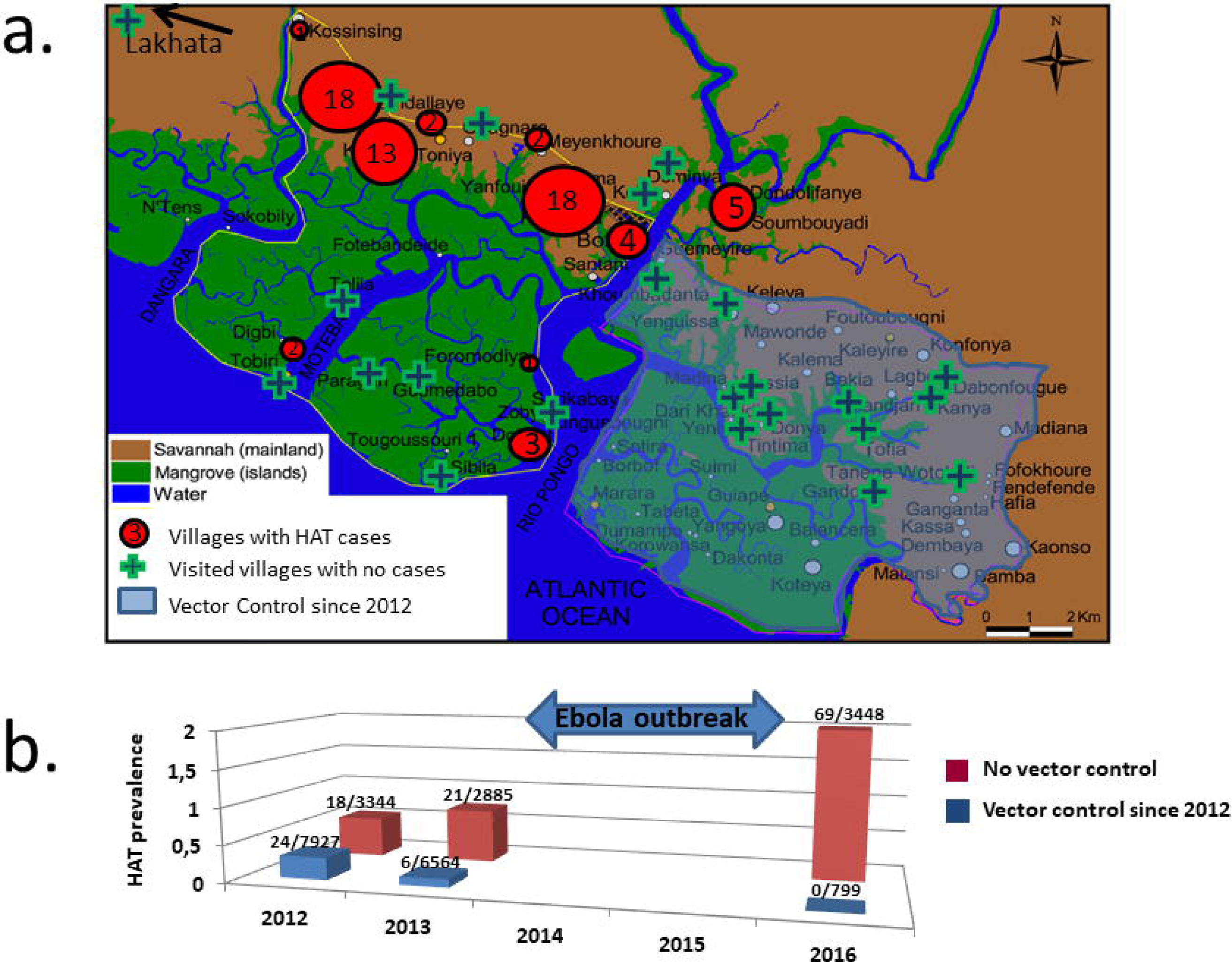
**a.** Geographic distribution of HAT cases diagnosed during two medical surveys led in the Boffa focus in 2016. **b.** Evolution of sleeping sickness prevalence in the Boffa focus assessed during active screening campaigns conducted in 2012, 2013 and 2016. The number of diagnosed HAT cases /number of persons screened is indicated above the histograms.

## Discussion and perspectives

A major lesson taken from the ebola outbreak is that medical care disruption may lead to quick HAT burst in areas where human are highly exposed to infective tsetse bites. In Boffa, epidemic levels were reached after only two years, ruining previous efforts to control the disease in areas with no vector control. This further indicates that albeit a control of disease prevalence can be reached by active screening campaigns, this strategy when implemented alone has poor effects on transmission. Tiny targets deployment in areas of human activities to reduce human exposure thus appears as a real preventive measure, remaining effective even in case of medical disruption. Once set-up, tiny targets remain active for a year and can be replaced by the communities themselves. We do believe that these observations are important to take into account when thinking tomorrow’s elimination strategies. In Guinea, tsetse control interventions are progressively extended to all HAT active foci, including the Western part of the Boffa focus but also the Dubreka and Forécariah foci and will help we hope keep in line with the 2020 elimination objective.

## Acknowledgments

We would like to thank all the health agents from the HAT National Control Program and the Boffa heath authorities as well as the villagers who actively participated in maintaining HAT control activities despite the challenge imposed by the ebola outbreak. This work was supported by the “Targeting Tsetse: a demonstration project” and the “Trypa-NO!” projects financed by the Bill & Melinda Gates foundation.

